# Parental state dynamically reshapes auditory processing of offspring vocalizations in zebra finches

**DOI:** 10.64898/2026.01.27.702116

**Authors:** Kristina O. Smiley, Felipe A. Cini, Luke Remage-Healey

## Abstract

Parental care is critical for offspring survival. For many species, including humans, auditory cues unique to dependent offspring, such as baby cries, elicit the necessary behavior from parents to care for young. Despite this, we know little about how the brain encodes auditory cues specific to offspring. Zebra finches are an excellent model to study this. Zebra finches are biparental, meaning both male and female parents raise the young and rely on auditory cues (begging calls) to elicit chick feeding responses (parental behavior). Begging calls also become individually identifiable by the parents, necessitating higher-order learning/ association neural processes. It is well established that the caudomedial nidopallium (NCM), a higher-order region of secondary pallial cortex (analogous to the mammalian secondary auditory cortex), is involved in the perception of complex auditory signals in birds. It is unknown, however, if/how NCM responds to offspring begging calls. To begin testing this, we used high density silicon probes to record single-unit *in vivo* electrophysiology activity in the NCM of adult parenting and non-parenting zebra finches exposed to playbacks of their own chick’s begging calls (parents only), novel age-matched begging calls (parents and non-parents), as well as novel adult male song and pure tone controls. Our results show that NCM neurons in parents respond more strongly (i.e., higher evoked firing rates) to both their own and novel chick begging calls and have higher baseline (spontaneous) firing rates in narrow spiking units relative to non-parents. Furthermore, NCM neurons in parenting females tend to show more selective responses towards their own chick begging calls, whereas those in males do not show similar levels of selectivity. These studies lay essential groundwork for future studies on how auditory responses can elicit parental behavior and how these responses may change over the chick-rearing period.

## Introduction

Parental care is an important adaptation observed across taxa to promote offspring survival. In many species, offspring use vocalizations or other auditory cues to signal distress, hunger, pain, coldness, and other needs (reviewed in^1–3)^. In turn, parents respond to these cues by providing the necessary care-giving behaviors toward offspring. The role of offspring vocalization cues has been well studied in the context of rodent maternal care (reviewed in^4^). For instance, in mice, dependent pups that are separated from the nest will emit ultrasonic vocalizations (USVs), which cause the mother to locate and retrieve the pup back to the nest^5^. In addition, mouse pups also emit another type of call, a wriggling call, which stimulates other maternal behaviors such as licking, changes in suckling position, and nest building^5^. In bilaterally deafened mothers, maternal behaviors are significantly reduced, causing slower growth and development of pups^5^. Therefore, exposure to these pup auditory cues is critical for maintaining the necessary level of postpartum maternal behavior required for successful pup rearing. In contrast, non-breeding or virgin rodents rarely respond behaviorally to pup cues^6^, and in fact, may even show aggressive or aversive behavior towards pups^7^. These observations suggest that changes in auditory processing may occur when animals enter a ‘parental state’ – allowing parents to respond appropriately to pup auditory cues, in ways that were not available in a non-breeding state.

Indeed, studies in rodents show that the primary auditory cortex undergoes significant plasticity following pregnancy and birth, which may alter the neural representation or perception of sounds, though the direction of this shift is not consistent across studies. While it is clear that that pup calls (USVs) generate strong, time-locked neural responses in the auditory cortex^8^, various studies have found that, compared to naïve females, mothers have either increased, decreased, or no change in evoked firing rates in response to pup USVs (for a thorough review and summary of studies, see^9^). In addition, spontaneous firing rates have been shown to both increase and decrease in mothers, compared to naïve virgins^9^. Thus, a general consensus of *how* auditory processing changes during motherhood has not yet been reached in rodents. In addition, there has been little work conducted on auditory plasticity outside of the murine primary auditory cortex (with the exception of the temporal association cortex (TeA) ^10^ and brainstem^11^), and virtually no studies to date have studied whether similar changes occur in male parents^1,9^, likely due to the fact very few rodent species are paternal. To better understand how parenting may shape auditory processing, we can make significant headway by studying species in which males and females can be studied.

Birds are an excellent model for the study of offspring vocalizations and the neural encoding of these calls. First, most birds are biparental^12^, meaning both males and females contribute to the raising of the young, allowing examination of sex-related effects on the response to offspring vocalizations. Second, like rodent pups, chicks use begging calls to elicit care-giving behaviors from adults. For birds, this is usually in the context of food begging. Food begging calls are generally thought to encode information about nutritional need, as spectral features of begging calls are modulated by hunger status and chick size^13–19^ (but see^2,3^ for more nuanced descriptions). Accordingly, parents adjust their amount of chick-directed feeding behavior in response to begging calls, with parents generally showing increased feeding rates towards nestlings that have been food deprived for longer^20–25^ (but see^26^ for an exception) and when additional playbacks of begging calls are played at the nest^20,23,27–30^. In contrast, surgically deafened parents do not sufficiently feed their offspring enough and abandon them prematurely, resulting in lower chick survival^31^. Other information such as chick identity, sibling order, position in the nest, predation risk, sex (of either the offspring or parent), age, and brood size may also be signaled in begging calls and affect parents’ responsiveness^3,32–38^. While the behavioral relevance of chick begging is clear, the neural mechanisms encoding begging calls, and how this encoding may change in parents, is not well understood in birds.

The auditory system has been thoroughly described in detail in birds, particularly in oscine passerines (songbirds) due to their pronounced ability to learn song^39^. Briefly, sounds are transduced from the cochlea to the brainstem via the auditory nerve and then auditory-driven activity travels through the midbrain and thalamus until reaching the auditory forebrain regions^40^. Here, activity reaches the pallium and is processed first in the primary thalmorecipient Field L (the primary auditory cortex analogue in birds) and then in the caudomedial nidopallium (NCM) and caudomedial mesopallium (CMM) – two secondary higher-order auditory regions in the avian brain^40^ (See Figure 1A for more details).

**Figure 1.**
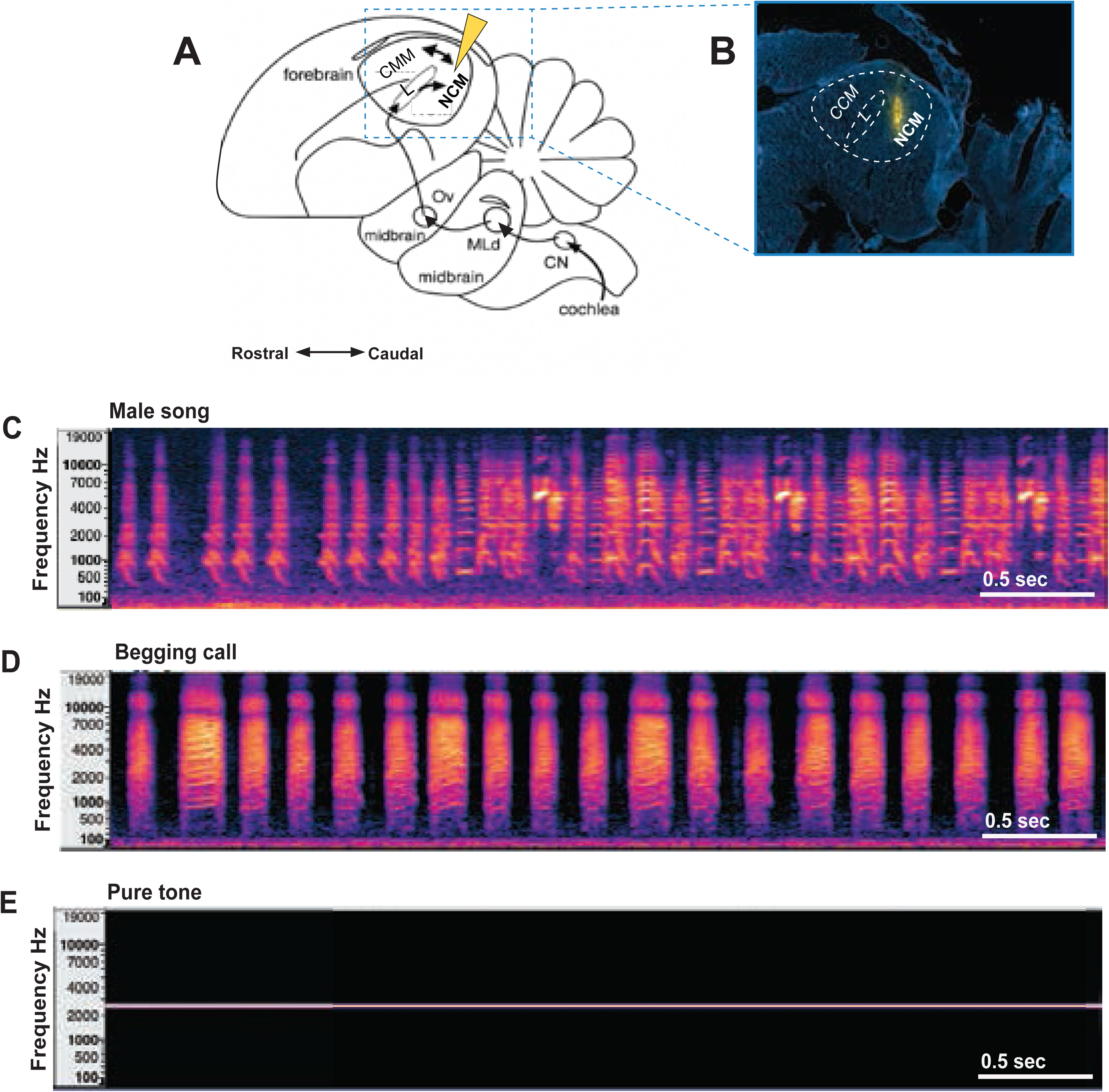
Recording site and auditory stimuli used. (A) Schematic of zebra finch brain auditory processing circuit in parasagittal view. The auditory nerve carries sound information into the brainstem, where it first reaches the cochlear nucleus (CN). Neurons from the CN send signals onward to the dorsal lateral mesencephalic nucleus (MLd) in the auditory midbrain. MLd then relays this information to the auditory thalamus, the nucleus ovoidalis (Ov). From Ov, the pathway continues to Field L (L), the primary thalamic target within the auditory forebrain. Field L in turn sends projections to the caudomedial nidopallium (NCM), which maintains two-way connections with the caudomedial mesopallium (CMM). Our probes were targeted at NCM (yellow arrow) for electrophysiological recordings (see methods). Blue dashed line represents area in shown in panel B. (B) Representative image of probe placement. Probes were dipped in fluorescent dye (yellow stain) to allow for posthumous site confirmation. Images were taken at 5X magnification. (C-E) Representative spectrograms of auditory stimuli used during the play back experiment. Each playback clip was 4 seconds long. Scale bar is 0.5 seconds (sec). (C) Representative adult male zebra finch stimuli used. (D) Representative begging call from a day 12 old chick. (E) Example of pure tone control stimulus used.

Within this pathway, we hypothesize that NCM is a critical region involved in the neural encoding of chick begging calls. One pinnacle of the auditory pallium (alongside CMM), NCM is involved in higher-order processing such sound categorization, memory, and recognition, and shows highly selective responses towards behaviorally relevant, conspecific, and familiar vocalizations^40–47^. For instance, in zebra finches (*Taeniopygia guttata*), NCM shows a greater response strength (i.e., evoked firing rate) and selective habituation towards conspecific songs, relative to heterospecific songs and pure tones, as evidenced by both immediate early gene studies and *in vivo* electrophysiology recordings (e.g., ^48–52)^. In addition, NCM is highly plastic, showing changes not only with experience-based song learning and memory^53–55^, but also in response to social interactions^56,57^, endocrine state^58^, breeding stage^59^, and seasonality^60,61^. Taken together, these findings suggest that the auditory processing mechanisms of NCM are adjusted to match the animal’s current social, physiological, and environmental state in birds.

Ample evidence demonstrates that the neural representation of a mate’s call or courtship song shifts depending on the breeding condition of the receiver^56,59,62^. Just as courtship signals become more salient during the breeding season, it is plausible that chick vocalizations become more salient when birds are in a ‘parental state’, reflecting a similar state-dependent tuning of the auditory system. While there is a vast literature on the role of NCM and other auditory processing regions in the context of courtship song learning and perception (much beyond what is reviewed here), in contrast, only one study to date has looked at the role of these regions in the context of begging calls^63^. In this study, Vidas-Guscic et al. found that offspring begging calls elicited stronger auditory responses (as measured by BOLD fMRI) than adult song motifs or tones in Field L and NCM of adult European starlings, regardless of season or sex^63^. Additionally, breeding-condition birds showed a selective seasonal increase in NCM responses to begging calls^63^, suggesting seasonal plasticity in processing socially relevant offspring signals. These findings further indicate that NCM is a promising target to investigate whether there are auditory processing differences in the neuronal response properties of parenting and non-parenting birds.

To dive further into the role of NCM in the encoding of chick calls, we sought to test whether NCM responses differed between parents (who are actively raising young) and non-parents (reproductively naïve individuals). To do this, we took an electrophysiological approach using adult zebra finches. Zebra finches are biparental, opportunistic and colonial breeders that originate from the arid regions of Australia but are easily kept in captivity and retain their naturalistic behaviors in laboratory settings^64^. Zebra finches can essentially reproduce anytime of the year (in the lab or the wild), as long as the heat and humidity levels are sufficient, and will lay consecutive clutches of 4-5 chicks for as long as breeding conditions remain favorable^64^. Zebra finch chicks use food begging displays that consist of both visual (mouth markings) and auditory (begging calls) components^30^. At very young ages begging displays are often silent or barely audible outside the nest. As chicks progress in their development, auditory begging calls become more prominent in their displays and progressively increase in loudness and intensity with age. By day 12 post-hatch, playbacks of begging calls alone are sufficient to stimulate parental feeding behavior^30^. Parents respond to begging calls by increasing their own eating first (of seeds), then shortly after will regurgitate this food to chicks. Male and female parents spend similar times feeding chicks throughout the day^37^.

A substantial point related to our exploration of the higher auditory pallium is that begging calls become individually identifiable by the parents, and parents show behavioral preferences for their own chick’s call, relative to other age-matched begging calls^38^. This suggests that higher-order learning/association neural processes are recruited to enable parents to recognize and discriminate among begging calls. In contrast, non-breeding zebra finches will not readily show care toward chicks^65^, suggesting that non-parents may have different auditory responses to begging calls. Here, we hypothesize that the activity of neurons in NCM reflects the encoding of begging calls in zebra finches. Specifically, we predict that parents will show differential processing (e.g., increased responsiveness and selective tuning) relative to non-parents, and that parents will respond stronger to their own chick, relative to an age-matched novel chick. Because males and females behaviorally respond similarly to begging calls^30^, we do not have *a priori* predictions on sex differences in NCM activity in the parenting context. This study is the first to measure neural activity responses using *in vivo* electrophysiology in zebra finches and will lay the essential groundwork for future studies to probe deeper into NCM’s role in auditory processing of begging calls and the mechanisms which modulate these calls during parenting.

## Methods

All procedures were approved by the Institutional Animal Care and Use Committee (IACUC) at the University of Massachusetts Amherst.

### Subjects

All subject birds were sourced from the Remage-Healey lab colony at the University of Massachusetts Amherst. Birds were kept on a 14:10 hr light:dark cycle, and group housed in mixed-sex aviaries with *ad lib* access to food, cuttlebone, and water. Diets were supplemented with egg food, fresh vegetables, and fresh millet weekly. The mean age of subjects at the time of the experiment was 438 ± 93 days. A total of 12 zebra finches were used (n=6 males and 6 females). Half of the subjects were parenting (n= 3 males, 3 females) and half of the subjects were non-parenting (n= 3 males, 3 females). Birds in the parenting condition were part of known pairs from the Remage-Healey colony. Exact reproductive histories of these birds were unknown but were presumed to have some reproductive experience prior to this study as these aviaries regularly had nestboxes on them which allowed for regular in-house breeding for the lab. Breeding status at the time of the experiment was monitored by nest observations and daily egg/chick counts inside the nest. The day the first chick hatched in the clutch was counted as “day 1 parenting”. Parenting birds were tested on “day 13 parenting”. Only one animal per breeding pair was used as a study subject, so that the other parent could remain with the chicks while the subject animal was being tested and after the study ended. This also meant that parenting subjects could remain statistically independent of each other as each subject came from a separate breeding pair. Each parenting nest used had an average of 4 ± 1 chicks in the nest at the time of testing (which is the average clutch size for zebra finches^64^). Non-parenting birds were age-matched to the birds in the parenting condition but had been kept in separate aviaries that never had nest boxes, and therefore, these individuals were known to be reproductively naïve. Sex of all subjects was determined by external plumage^64^.

### Experimental Design

All subjects underwent craniotomy surgery 1-2 days prior to testing (i.e., when *in vivo* electrophysiology recordings were made). Parenting birds were removed from the colony room on “day 12 of parenting” to undergo surgery. During this time, the nest box belonging to that individual was also removed and brought into a sound attenuated chamber during the duration of the surgery (average 2 hours). Following surgery, the begging call from the eldest chick (i.e., the chick that was 12 days old) was recorded for several minutes to generate stimuli for ‘own chick begging call’ (see details below). Once the call recording was complete, the nestbox was promptly replaced back in its original location in the colony room so that the other parent could continue to care for the chicks. Parenting subjects were kept in single-housed recovery cages, alongside several other birds in single-house cages, in a separate room away from the colony and their nest box following surgery and for the remainder of the experiment. Non-parents underwent similar procedures under a similar timeframe but did not have any chicks to record.

One to two days following the surgery, electrophysiological recordings were collected from NCM while birds heard auditory stimuli (see details below). After all recordings (1-2 days) were complete, subjects were perfused, brains were collected and sectioned, and electrode placement was verified (details below).

### Craniotomy Surgery

Craniotomy procedures were followed as previously described^66^. Briefly, birds were fasted for 30 minutes prior to surgery and then anesthetized with 2% isoflurane in 2 L/min O_2_, fixed to a custom stereotax (Herb Adams Engineering) equipped with a heating pad (DC Neurocraft) at a 45° head angle, and then maintained on ∼1.5% isoflurane in 1 L/min O_2_ for the duration of the surgery. Lidocaine (2% in EtOH, vol = 0.02 ml) was injected subcutaneously in the skin covering the top of the bird’s head prior to making an incision to expose the skull. Skull markings were etched over NCM in both hemispheres (0.9 mm rostral and 0.7 mm lateral from the bifurcation of the mid-sagittal sinus), and large craniotomies over NCM were made and the meninges were resected. Custom-made metal headposts were lowered on the top of the beak and secured with dental cement. A small craniotomy was also made in the skull over the cerebellum for the implantation of a silver ground wire. Craniotomies were sealed with Kwik-Cast (World Precision Instruments), and birds were allowed to recover from anesthesia for several minutes before being transferred to individually housed recovery cages. Birds were given meloxicam (0.3 mg/ml, vol=15 μl) before the start of surgery and the following morning (within 24 hours after surgery) for pain management.

### *In vivo* awake, head-fixed electrophysiology recordings

Recording procedures were followed as previously described^66^. Briefly, on the day of the recording, birds were moved into a sound attenuation booth (Industrial Acoustics) and comfortably restrained in a cloth jacket and plastic tube and head fixed on the recording rig. The Kwik-Cast was removed from the craniotomy over one of the hemispheres and silicon oil was placed over the exposed surface of the brain. Sixty-four or 128 channel silicon probes (Diagnostic Biochips/Cambridge NeuroTech) were fixed to a micromanipulator and positioned over the subject’s head. Before insertion in the brain, electrodes were dipped in DiI-594 (6.25% in 100% ethanol, ThermoFisher Scientific) for posthumous site confirmation. Probes were lowered into NCM 1.5-2.0 mm ventral from the brain surface. After finding a site with characteristic NCM baseline and sound-evoked activity across the recording channels (387.5 μm in length), the probe was allowed to stabilize in the brain for 30-60 minutes.

Recordings were made while animals listened to auditory stimuli. Three different recording sessions per hemisphere were conducted, each using a unique set of auditory stimuli. Each stimulus set contained one novel male song, one novel chick begging call, one own chick begging call (parents only), and one pure tone control (see below for details). The speaker was positioned ∼30 cm from the animal during playbacks, and sound level was amplified to 65 dB as measured by a sound level meter at the animal’s position (RadioShack). Each sound was played for 4 seconds, repeated 15 times in pseudorandom order, and the inter-stimulus presentation interval was randomized to be between 2 and 10 seconds long. Each playback trial duration lasted ∼11 minutes.

Recordings were sampled at 30 kHz using Open Ephys software^67^. An Arduino Uno (Arduino) was connected to the recording computer to deliver TTL pulses to the evaluation board’s DAC channel bracketing the beginning and end of the audio stimuli to optimize detection during analysis. Audio playback and TTL pulses were controlled by a custom-made MATLAB (MathWorks) script which also controlled the Arduino and sent a copy of the audio analog signal to the evaluation board ADC channel.

Recordings from each hemisphere were performed on the same day if possible (alternating starting with the left and right hemisphere between subjects). If recording both hemispheres on one day was not possible, then subjects were kept in their recovery housing cage overnight and recorded again on the following day. Birds were recorded for no longer than 4 hours per day.

### Brain collection, sectioning, and imaging (electrode verification)

At the end of the experiment, subjects were euthanized by an overdose of isoflurane vapor and perfused transcranially with phosphate-buffered (PB) solution followed by 4% paraformaldehyde in PB. Brains were removed and post-fixed in 4% paraformaldehyde overnight before they were transferred to 30% sucrose in PB for 2-3 days (or until they were fully sunk). Brains were then embedded in OCT inside small plastic molds and then frozen at −20 °C until they were sectioned.

Brains were sectioned on a cryostat into 40 um sections and mounted serially onto SuperFrost™ slides (Thermo Scientific™). Slides were air dried in the refrigerator before being rinsed in PB and then cover slipped using ProLong™ Gold Antifade Mountant containing DAPI (Invitrogen™). Slides cured overnight and then were imaged on a Zeiss ApoTome.2 microscope equipped with an Axiocam 105 color camera, using Zeiss Zen 3.7 software (refer to Figure 1B for an example verification of electrode placement in NCM).

### Auditory Stimuli

Three separate sets of auditory stimuli were created, each containing one novel male song, one novel chick begging call, one own chick begging call (parents only), and one pure tone control. All stimuli were saved as 16-bit .wav files and mean amplitude-normalized to 65 dB using Audacity® (version 3.4.2., https://sourceforge.net/projects/audacity/).

Three separate male song recordings were randomly chosen from a data set provided in^41^ so that they were unfamiliar to our subjects. Each song was trimmed to 4 seconds, containing introductory notes and at least 2 motifs (see Figure 1C for an exemplar spectrogram), and then was randomly assigned to a stimulus set. Novel male songs were used as ‘positive control’ as NCM reliably displays robust responses to male song in both male and female zebra finches (e.g., ^40–43)^.

‘Novel begging calls’ were selected from a library of begging calls that were collected from separate nests in the Remage-Healey breeding colony several months prior to this study, and therefore, were assumed to be novel to all birds used in this study. Three separate age-matched chicks (all 12 days post-hatch) were chosen at random to use and then randomly assigned to a stimulus set. Chick begging calls at this age contain small ‘begging bouts’ that last ∼0.1 s with ∼0.1 s silence in between bouts. All begging call stimuli were trimmed to 4 seconds, with each trimmed section containing ∼20 bouts of begging. Areas within the recording that were chosen as sections to trim aimed to include only full bouts of begging, so that they were more ‘natural sounding’ (i.e. – not cut off in the middle of a begging bout)-see Figure 1D for an exemplar spectrogram.

For parenting subjects, recordings were made from their eldest chick (age 12 days post-hatch) on the day they underwent craniotomy surgery for ‘own chick begging call’ stimulus. Zebra finch chicks hatch sequentially (1 day apart), therefore, only one chick per nest would be aged 12 days post-hatch at the time of recording. As we wanted to keep all chick stimuli age-matched between the novel and own chick begging call groups, we only used the eldest chick from each parent’s nest (on day 12 post-hatch) for ‘own chick begging call’. However, we used three different trimmed sections from the own chick recording (see below), and each trimmed section was randomly assigned to a stimulus set.

Finally, 3 different pure tones were used (2000, 2200, and 2600 Hz) and were randomly assigned to a stimulus set (see Figure 1E). These served as ‘negative controls’ as it is well established that NCM shows lower activity responses to pure tones (e.g., ^40–43)^.

### Begging call recordings

Begging calls were recorded inside a sound attenuated chamber equipped with a lave-ier microphone. Chicks were separated from their home cage inside their nestbox for ∼2 hours prior to recording. At the time of recording, one chick was taken out of the nest and placed in a small paper-lined bowl with the microphone attached to the side. As begging calls can be elicited by movements to the mouth, a ‘fake adult beak’ (a piece of lab tape wrapped around the end of a pen) was used to tap the chick’s beak or tongue to elicit begging calls. Each chick was recorded for ∼ 2 minutes. Sounds were recorded into the program Sound Analysis Pro (https://soundanalysispro.com/) and saved as .wav files.

### Data analysis

Data analysis was conducted as previously described^66^. Briefly, sound playback timestamps were detected using a custom-made audio convolution algorithm in MATLAB. Recordings were highpass filtered at 300 Hz and common median filtered in MATLAB (MathWorks). Single-unit sorting was done with Kilosort4^68^ and then manually curated in Phy (https://github.com/cortex-lab/phy). Only well-isolated units were used (high signal-to-noise ratio; low violation of refractory period (the number of interspike intervals within a 1 ms refractory period was 0.25% median in our units), and low contamination with other units (segregation in waveform PCA space). Cells with spike widths ≤0.4 ms were classified as narrow spiking, while cells with spike widths >0.4 ms were classified as broad spiking^66^; these classifications correspond to inhibitory and excitatory neurons in zebra finch NCM^69^. All further data processing was conducted in Python.

### Statistics and data visualization

All statistical analyses and plotting were performed using libraries in Python. Data were fitted with Gaussian distributions and determined to be normally distributed as assessed by visual inspection of Kernel Density Estimate histogram plots of model residuals, Quantile-Quantile plots, and Residuals vs. Fitted plots.

### Evoked NCM responses across treatment groups

To measure differences in evoked responses across treatment groups (Figure 2), Z scores were calculated by the formula below, where S and B are the stimulus and spontaneous baseline firing rates across stimulus trials, respectively.

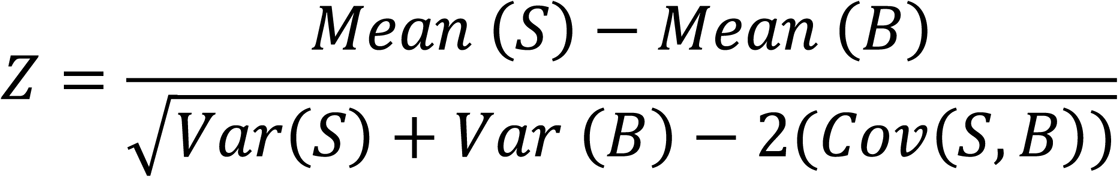

**Figure 2.**
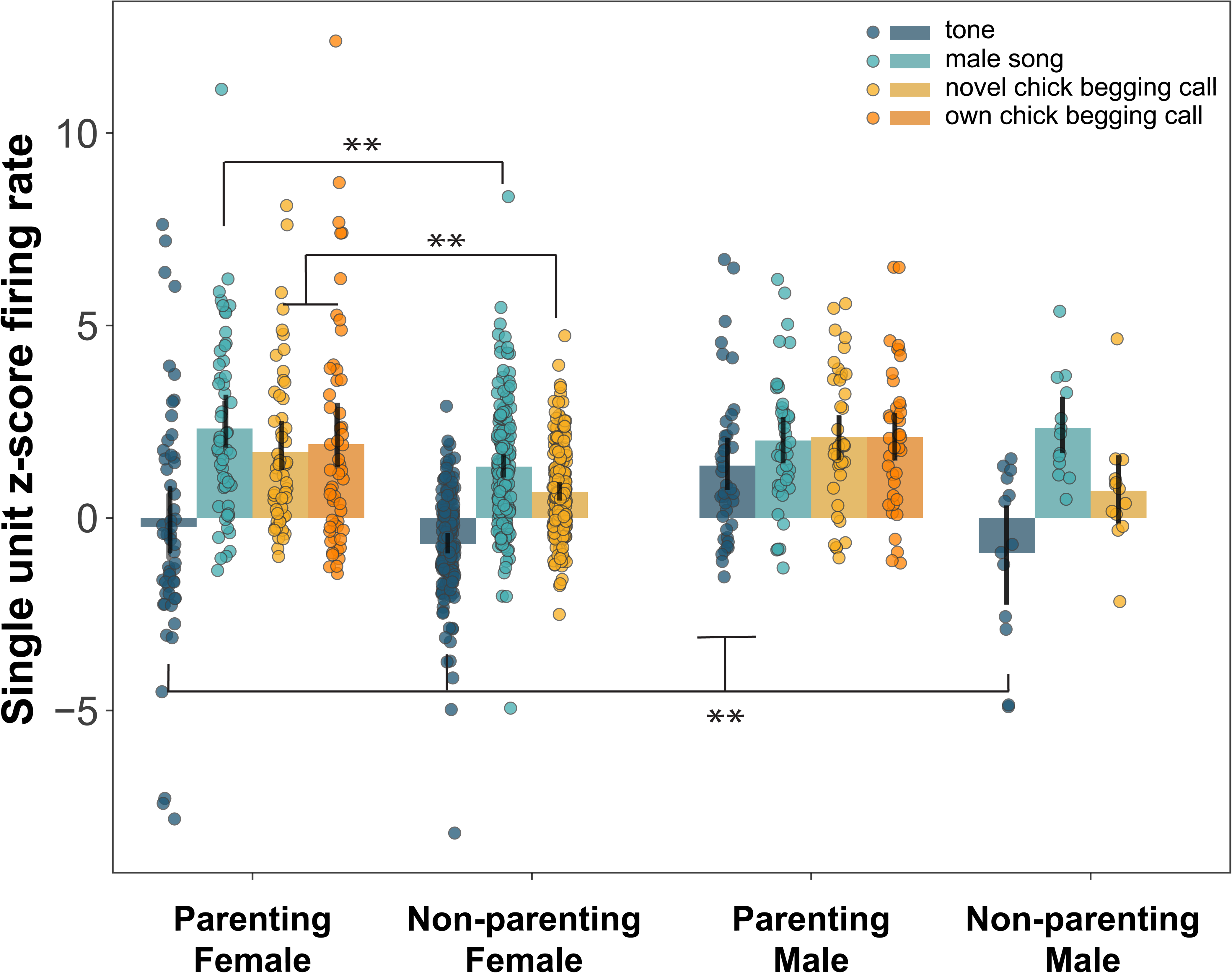
Stimulus-evoked firing rates in NCM are increased in parenting females and males. Shaded bars depict mean firing rates to each stimulus, across experimental groups. Black lines indicate standard error of the mean. Filled circles represent single data points (i.e., single units measured). Neurons in parenting females had significantly higher firing rates to song and to both novel and own chick begging calls, relative to non-parenting females. Neurons in parenting males, while also increasing firing rates towards both songs and begging calls, showed significantly higher responses to tone, relative to all other groups tests. Note that only parenting females and males heard their own chick’s begging calls, as non-parenting birds did not have chicks at the time of recording. **p<0.01. For a complete report of all pairwise post-hoc tests, see Table S1.

We then ran a 4-way ANOVA with parenting status (parenting or non-parenting), cell type (broad or narrow spiking neurons), stimulus type (tone, male song, novel begging call, own begging call), and sex (male or female) as the main factors and evoked response at the outcome variable. Post-hoc tests were conducted with Tukey HSD tests. Note, we were unable to record or obtain units from both hemispheres for every subject and therefore were unable to include hemisphere as a factor.

### Selectivity measures

Selectivity distributions were generated by plotting the percentage of stimuli a single unit responded to relative to the density of neurons (i.e., percentage of neurons) which respond that a given percentage of stimuli for parenting males and parenting females (Figure 4A) and broad and narrow spiking neurons (Figure 5B). These distributions were compared using a Kolmogorov-Smirnov test.

To test whether there was a preference in the stimulus types parenting females and male were responding to, we calculated d-prime (d’) for each social stimulus (i.e., novel male song, novel chick begging call, and own begging call, using tone as reference), using the formula below where μ_1_, μ_2_= mean responses to each stimulus and σ_pooled_ = pooled standard deviation of the two response distributions. Differences in d’ were compared using ANOVA comparing effects of sex, stimulus, and sex × stimulus, followed by Tukey post-hoc tests (Figure 4B).

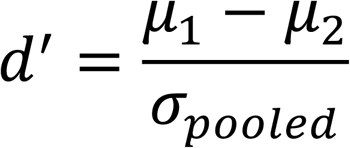

### Baseline firing rates

Spontaneous baseline firing rates were averaged during the 2 seconds prior to the onset presentation of each stimulus. The average baseline firing rates for broad and narrow spiking units were compared between parenting groups and sex using an ANOVA with Tukey post-hoc tests (Figure 5A).

## Results

In total, we recorded 283 single units (n=148 broad spiking units; n=135 narrow spiking units) from 8 different animals: Parenting males (n=2), Parenting females (n=2), Non-parenting males (n=1), Non-parenting females (n=2).

### Parents show greater evoked responses in NCM compared to non-parenting birds

We first examined the effects of sex (male or female), stimulus (own begging call, novel begging call, male song, or tone), parenting status (parental or non-parental), and cell type (narrow spiking or broad spiking neuron) on evoked NCM responses (i.e., firing rate). Using an ANOVA, we found a significant sex × stimulus × parenting status interaction (F(3, 910) = 3.42, p<0.01). Tukey post hoc tests revealed that neurons in parenting females showed a higher response to both their own (p<0.001) and novel begging calls (p=0.02), compared to those in non-parenting females exposed to novel begging calls (Figure 2). Female parents also had neurons with a greater response to male song compared to non-breeding females (p=0.03). For both parenting and non-parenting females, neurons had a significantly lower response to tone compared to all other stimuli (all p<0.05). On the contrary, neurons in parenting males showed heightened responses to all stimuli such that that responses to tone were no different from the other stimuli presented (all p>0.05). In addition, neurons in parenting males had a significantly higher tone response compared to the tone response of parenting and non-parenting females and non-parenting males (all p<0.01; Figure 2). For a complete reporting of all post-hoc tests, refer to Supplemental Table S1. For exemplar single unit traces from each treatment group to each auditory stimulus, refer to Figure 3.

**Figure 3.**
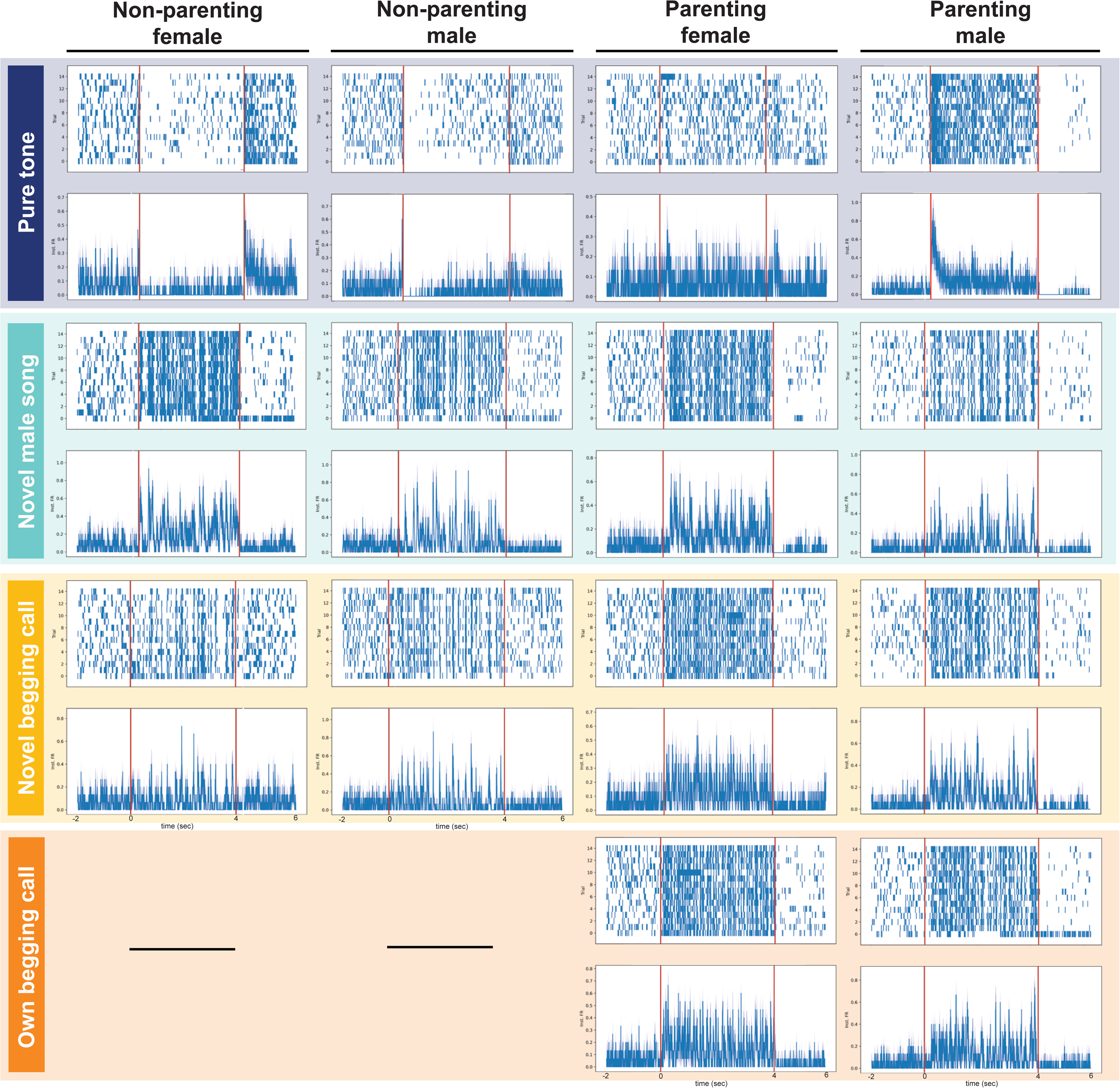
Single unit exemplar raster plots and average instantaneous firing rates per stimulus, across groups. Each column represents one of four treatment groups of animals: Non-parenting females, non-parenting males, parenting females, and parentings males. Each row is colored by stimulus type: Pure tone (blue), novel male song (turquoise), novel begging call (yellow), and own begging call (orange). Note that only parenting females and males heard their own chick’s begging calls, as non-parenting birds did not have chicks at the time of recording. For each figure, the top panel depicts the spiking activity over the 15 trials (X-axis) the stimulus was played. The bottom panel depicts the average instantaneous firing rate (Inst. FR; X-axis) across the trial. For both panels, the Y-axis is time in seconds (sec). Note each graph displays the 2 seconds prior to the onset of the auditory stimulus (time 0), which was played for 4 seconds, as well as 2 seconds after the offset of the stimulus. The onset and offset of the auditory stimulus are marked by the red lines in each panel. These data correspond with the significant differences found in evoked firing rates displayed in Figure 2.

### Parenting females are more selective in their auditory responses than parenting males

Although neurons in parenting females and males had similar evoked response strength for begging calls and male song, females appeared to be differentiating their response somewhat between stimuli, whereas males responded to all stimuli similarity (Figure 2). To test whether there were in fact differences in the selectivity to auditory stimuli, we compared the selectivity distributions of parenting females and parenting males using a Kolmogorov-Smirnov test. Indeed, females showed a greater density of neurons that are more selective (i.e., respond to a lower percentage of stimuli), relative to males (D = 0.27, p = 0.003; Figure 4A). When selectivity between sexes was compared using d’, neurons in females were also nearly significantly found to be more selective than males (F(1, 60) = 3.62, p=0.06; Figure 4B). When we tested for differences in selectivity towards different social stimuli type within each sex using d’, we found a trending, but non-significant difference in selectivity by stimulus type in parenting females (F(2, 60) = 2.86, p=0.07; Figure 4B), but not males. To visualize this, we plotted each single unit by time and response selectivity to each social stimulus. Overall, neurons in females tend to show more selective responses towards their own chick calls, relative to novel chick calls (Figure 4C-E), and male song (Supplemental Figure), whereas males do not show a similar level of selectivity (Figure 4F-H; Supplemental Figure).

**Figure 4.**
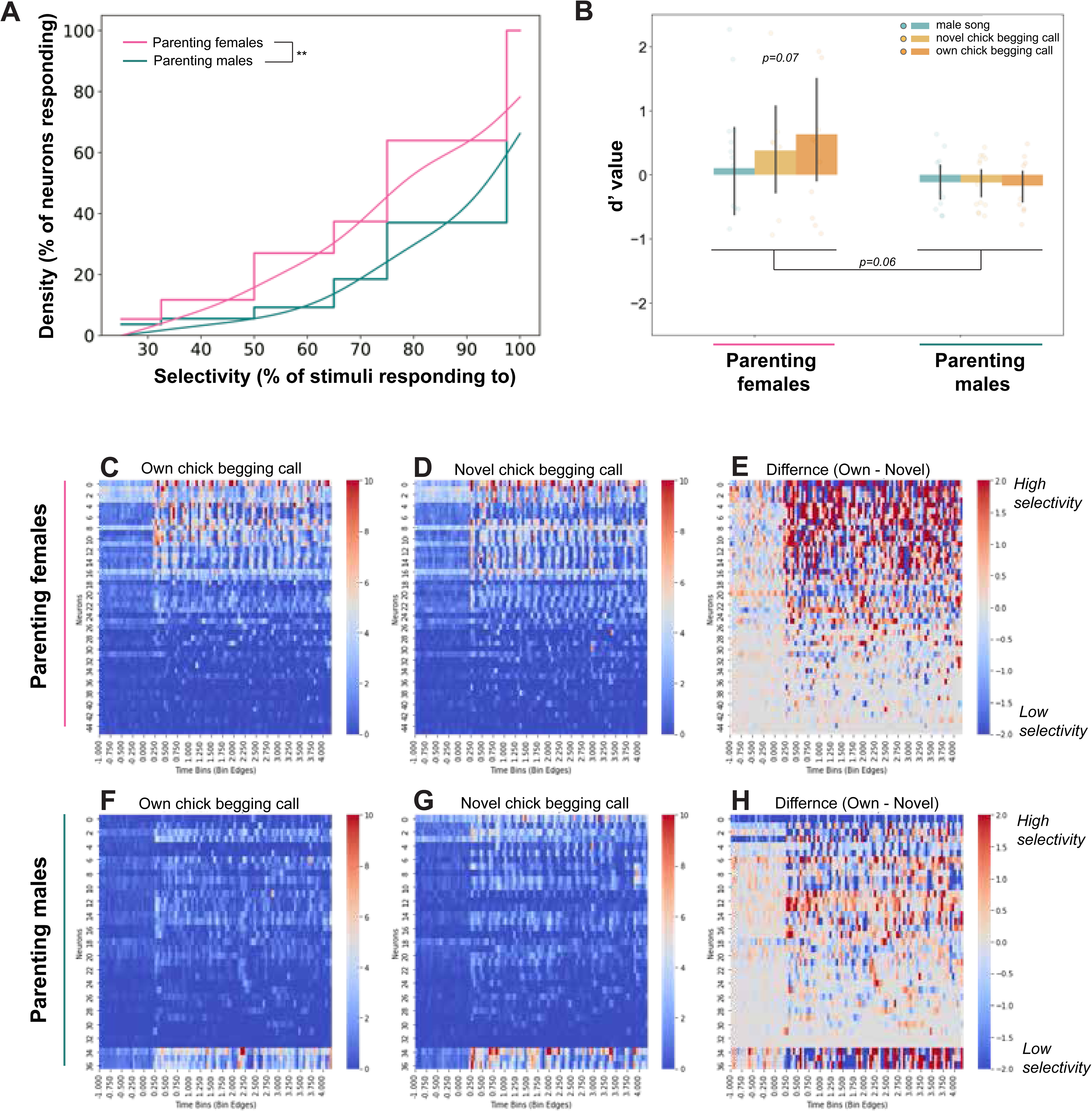
Parenting females are more selective in their auditory responses, compared to parenting males. (A) Here selectivity is plotted as the percentage of stimuli a single unit responds to (X-axis) relative to the density of neurons (i.e., percentage of neurons) which respond that a given percentage of stimuli (Y-axis). Overall, parenting females show a greater density of neurons that are more selective (i.e., respond to a lower percentage of stimuli). **p<0.01, Kolmogorov-Smirnov test. (B) In agreement with panel (A), d’ values also indicate that overall, parenting females were trending to be more selective than parenting males (p=0.06). Of the social stimuli presented (male song, novel and own chick begging calls), d’ values indicate that parenting females trended towards showing the strongest selectivity for their own chick (p=0.07), whereas males showed little selectivity towards any stimulus. (C) Visual depiction of single unit selectivity for parenting females and males for their own vs novel chick begging calls, across the trial period. Each Y-axis row represents a single unit recording. The X-axis are time bins (50 ms) divided across the duration of the playback trial. Plotted colors are the average firing rates for each unit at a particular time bin. Warmer colors (reds) depict more selective responses, whereas cooler colors (blues) depict more unselective responses. White/neutral colors indicate no selectivity toward either stimulus. The last plot for both parenting females and males is the difference in response for own – novel chick begging calls, with warmer colors (reds) depict more selective responses towards their own chick’s begging call. Note that females show a greater number of units that show stronger selectivity towards their own chick relative to males listening to their own chick.

### Spontaneous baseline firing rates of narrow spiking neurons are higher in parents compared to non-parents

To test whether baseline firing rates differed between treatment groups we compared the average baseline firing rates (2 seconds before the presentation of the stimulus) between parents and non-parents. Using an ANOVA, we found a significant sex × cell type interaction (F(1, 934) = 2.67, p<0.01). Tukey post hoc tests revealed that parents had higher baseline firing rate in narrow spiking units, relative to narrow spiking units in non-parents (p<0.01), but there was no difference in baseline broad spiking neurons (p=0.82; Figure 5A). Consistent with other previous literature^70^, narrow spiking neurons had a higher baseline firing rate compared to broad spiking neurons across all groups (Figure 5A).

**Figure 5.**
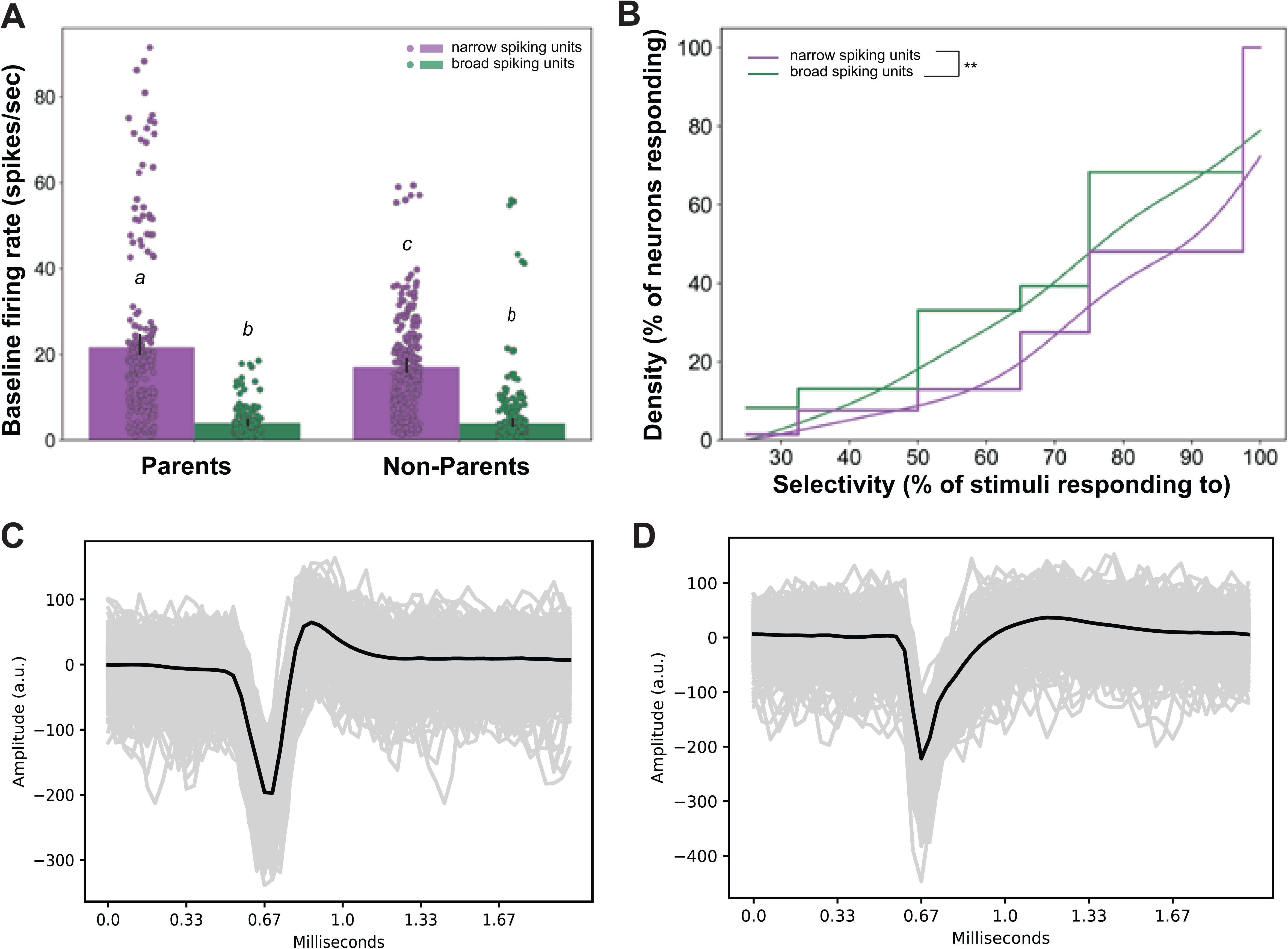
Parents have higher baseline firing rates in narrow spiking neurons, relative to non-parents. (A) Each dot represents a single unit; bar plots are average baseline firing rates (2 seconds before the presentation of the stimulus) ± standard error of the mean. Differing letters above the bar plots indicate significant pairwise differences (Tukey post hoc test, p<0.05). (B) Here selectivity is plotted as the percentage of stimuli a single unit responds to (X-axis) relative to the density of neurons (i.e., percentage of neurons) which respond that a given percentage of stimuli (Y-axis). Overall, broad spiking neurons are more selective than narrow spiking neurons. **p<0.01, Kolmogorov-Smirnov test. (C,D) Exemplar waveforms of narrow spiking (C) and broad spiking (D) neurons.

### Broad spiking neurons are more selective than narrow spiking units

To test whether there were differences in cell-type specificity, we compared the selectivity distributions of broad and narrow spiking neurons using a Kolmogorov-Smirnov test. Overall, broad spiking neurons show a greater density of neurons that are more selective (i.e., respond to a lower percentage of stimuli), relative to narrow spiking neurons (D = 0.201, p = 0.006; Figure 5B).

## Discussion

### Overview of results

We tested whether the activity of neurons in NCM reflects the encoding of begging calls in zebra finches. Using *in vivo* extracellular recordings, we found three main results. First, relative to reproductively naïve birds, neurons in parenting birds showed enhanced evoked responses in NCM to social sounds—both chicks begging calls (own and novel) and novel male song. Interestingly, we found that parenting females tended to be more selective in their NCM responses, favoring their own chick’s begging calls, over other social sounds, while neurons in males displayed a broader elevation of responsiveness, including tones, and displayed lower response selectivity than parenting females. Lastly, spontaneous (baseline) firing rates were higher in narrow-spiking (putative inhibitory) units in parents than in non-parents, while broad-spiking (putative excitatory) units were more selective overall than narrow-spiking units. Together, these results indicate state-dependent modulation of auditory processing in NCM during parenting, with sex-specific response profiles.

### Parents show greater evoked responses in NCM compared to non-parenting birds

In support of our hypothesis, we found that parents exhibited stronger NCM responses toward chick begging calls than non-parents. This finding suggests that entering a “parental state” may alter auditory processing to enhance sensitivity to behaviorally relevant signals from dependent offspring. Comparable state-dependent modulation of auditory responsiveness has been well-documented in the context of courtship communication. In many songbird species, breeding or pair-bonded individuals show heightened neural and behavioral responses to mate calls and/or male courtship song relative to unfamiliar calls and non-courtship songs^56,59,62^.

This increase in responsiveness may be driven by the endocrine shifts that occur when animals transition into breeding state. For example, in female white-throated sparrows (*Zonotrichia albicollis*), immediate early gene responses in NCM to male courtship song is enhanced when females are treated with high-levels of estradiol, a condition that mimics the endocrine breeding state, but not under low-estradiol conditions, which mimics a non-breeding state^70^. By analogy, chick vocalizations may gain more salience when birds transition into a parental endocrine state, reflecting a parallel form of hormone-dependent tuning of auditory circuits. Indeed, parenting is accompanied by marked shifts in circulating hormones, including elevated estrogen and progesterone during egg laying and elevated prolactin and mesotocin (avian homologue of oxytocin) during post-hatch care, which together orchestrate parental behaviors such as brooding and provisioning^71^. These hormones may also influence sensory processing. NCM neurons are known to express steroid hormone receptors, and much work has demonstrated direct estrogenic modulation of auditory responses to song (reviewed in^62,72^).

Thus, it is plausible that the endocrine changes associated with the onset of parenting could act directly on NCM neurons to increase responsiveness to chick calls. Although this hypothesis has yet to be directly tested in birds, there is some support from a study in mice which showed that oxytocin acting on auditory-cortex neurons enhances maternal responsiveness to pup vocalizations^73^. Hormonal effects may also operate indirectly through other neuromodulatory systems that project to NCM. For example, dopaminergic input from the ventral tegmental area or noradrenergic input from the locus coeruleus, both of which are also hormone-sensitive in songbirds^72^, could also modify responsiveness to behaviorally salient sounds. Future studies will be required to determine whether similar neuroendocrine mechanisms shape auditory processing during avian parental care.

Another, though not necessarily mutually exclusive, explanation is that the heightened responsiveness in parents reflects experience-dependent changes—specifically increased exposure, familiarity, learning opportunities, and potentially recognition of chick begging calls. Parents in our study were both reproductively experienced and had been actively caring for chicks for nearly two weeks before neural recording, giving them substantial exposure to chick calls. In contrast, non-breeding birds were reproductively naïve and had no prior adult experience with chick vocalizations. One possibility is that increased experience—or simply the greater cumulative exposure to chick vocalizations—refines auditory sensitivity through repeated activation of relevant neural circuits. Beyond sheer exposure, sustained interaction with their own brood could promote familiarity, strengthening auditory memory traces that bias attention toward these calls. Consistent with this, adult female zebra finches show enhanced NCM responses to familiar songs, including both their mate’s and other familiar males’ songs, as well as heightened selectivity for these signals^56,57,59^. These findings demonstrate that auditory familiarity reliably increases NCM responsiveness, suggesting a parallel mechanism may exist for parents that are repeatedly hearing their chicks’ calls. Whether such effects persist across breeding experiences or depend on current hormonal/behavioral state remains to be determined; testing this in non-breeding, but reproductively experienced adults would help disentangle these possibilities.

A related, but distinct, mechanism involves learning opportunities created during active parental care. For parents, chick begging calls are repeatedly paired with provisioning, assessment of need, and other meaningful parental outcomes. Song-learning studies show that pairing conspecific song playback with a reinforcing stimulus induces long-term changes in NCM responsiveness^74–76^, illustrating that associative learning can shape auditory processing. Such reinforcement-linked exposure could similarly enhance perceptual tuning to chick vocalizations during parenting. At a higher representational level, parents may also develop recognition abilities—distinguishing their own chicks from others—through experience with individualized begging calls. Zebra finches are capable of individual recognition of chicks based on call structure^38^, and NCM plays a role in both memory and individual vocal identity (reviewed in^77^). Thus, increased experience, familiarity, learning, and recognition may act together to enhance parental responsiveness to chick calls. These experience-dependent mechanisms could operate independently of, or in concert with, hormonally driven changes in auditory processing described above.

### Parents show higher baseline firing in narrow-spiking neurons, relative to non-parents

In addition to increased response strengths, both male and female parents also showed higher baseline firing rates in narrow spiking neurons (putative inhibitory interneurons). This could reflect a basic alteration in intrinsic properties via changes in ion channel conductances, such as those observed in songbird neurons recently^78,79^, and which could be hormone- and/or experience-dependent. Evidence from rodent studies have shown that inhibitory interneurons—especially parvalbumin-positive interneurons—are a primary site of experience-dependent plasticity in the auditory cortex during maternal care (reviewed in^9^). Compared to naïve virgin female mice, maternal mice exhibit inhibitory responses to pup calls that are earlier, longer-lasting, stronger, and more stereotyped, indicating a major reorganization of inhibitory processing with parental state (reviewed in^9^). In songbirds, song-evoked responses rely on GABA-mediated inhibition in NCM and inhibitory modulation of selectivity for different song components is necessary for the encoding of learned song categories^80^. While our data do not directly test whether inhibitory firing is directly related to changes in auditory processing of begging calls, it does suggest an overall change in inhibitory tone across the NCM of both female and male parents. This shift may alter the balance of excitation and inhibition in auditory circuits, possibly fine-tuning NCM’s responsiveness to auditory stimuli during parenting. This shift may be a result of hormonal action that coincides with the onset of parenting. In support of this hypothesis, Marlin et al. found that oxytocin (which increases with maternal care) acting in the auditory cortex of mice balanced the magnitude and timing of inhibition, leading to increased discrimination of pup calls and enhanced maternal behavior^73^. Whether such a phenomenon occurs in birds has not been tested but is certainly an intriguing possibility. If similar hormonal effects were to be found in the auditory processing areas of birds, this could unmask conserved mechanisms by which parents are able to ‘hone in’ on offspring stimuli which allows them to appropriately adjust their parental behavior in order to meet their offspring’s needs.

### Lack of neural discrimination between own and novel chick begging calls in parents

Despite previous evidence that zebra finches can behaviorally discriminate and recognize their own chicks based on begging calls^38^, contrary to our prediction, we did not observe any difference in NCM response strength between parents hearing their own versus an age-matched novel chick. Levréro et al.^38^ showed that by day 11 post-hatch, chick calls already contain individually identifiable acoustic features, suggesting that parents should, in principle, be able to distinguish between their own and unfamiliar offspring. We tested parents slightly later, at day 13 post-hatch, using playbacks of calls recorded from their eldest chick shortly after their last interaction prior to testing (see Methods). Nonetheless, we found no evidence of discrimination at the level of overall NCM response amplitude. Several factors may account for this absence of difference.

First, although chick calls are acoustically individualized by day 11, parents may require additional time or reinforcement to form and consolidate the learned association between specific call features and their own offspring. Behavioral studies demonstrating reliable offspring recognition in zebra finches typically involve older, post-fledgling-stage chicks^38,81^, suggesting that auditory discrimination may emerge later in development. Thus, testing parents when chicks were slightly older might have revealed stronger own-chick selectivity.

Second, discrimination between familiar and unfamiliar vocalizations might occur at another stage of the auditory processing hierarchy rather than being expressed in overall NCM response magnitude. Although NCM is known to differentiate between familiar and unfamiliar conspecific songs or calls (e.g.,^51,57,82^), subtle individual-level distinctions may instead be represented in finer-scale neural coding or in other downstream regions involved in recognition or behavioral decision-making.

Third, being in a parental state may broadly heighten responsiveness to any chick-related vocalizations that are behaviorally relevant, irrespective of relatedness. Indeed, NCM has been shown to recognize “categories” of natural sounds^83,84^ and therefore novel and own chicks may be grouped into one “begging call category”, thereby eliciting similar neural responses. One reason this may work is that many birds which raise altricial young rely on location or contextual cues to identify their young – and may not necessarily have/use individual identity for each chick. In zebra finches, which continue to provision fledglings outside the nest after fledging, selective recognition of individual begging calls may only become necessary—and therefore only expressed—closer to fledging age. Future experiments could test this hypothesis by comparing neural responses to own and novel chick calls across chick developmental stages.

Finally, although we did not detect population-level differences in response strength, single-unit analyses revealed a trend toward greater selectivity in females, with more neurons showing preferential responses to their own chick’s calls (Figure 4). This suggests that some degree of individual discrimination may already be emerging at the level of single neurons, particularly in females, even if it is not yet reflected in the mean NCM response amplitude.

### Sex differences in parents’ auditory responses towards begging calls

Although all parents showed enhanced responsiveness to chick vocalizations, males and females differed subtly in how these responses were expressed in NCM. Females exhibited greater single-unit selectivity toward their own chick’s calls relative to novel chick calls and male song (Figures 4, S1), suggesting some degree of emerging individual discrimination at the neuronal level. In contrast, males showed uniformly elevated responses across all tested stimuli—including tone—and exhibited little evidence of selective single-unit tuning. These findings were surprising for two reasons. First, pure tones typically elicit weak NCM responses in zebra finches, making the male-specific enhancement to tones comparable to those elicited by song and chick calls unexpected. Second, zebra finch males and females generally provide similar forms and amounts of parental care^64^, so diverging neural response patterns were not necessarily anticipated given their broadly shared behavioral roles.

Although speculative, one possibility is that these results reflect a case of sex convergence, in which similar behavioral phenotypes are supported by different underlying neural mechanisms^85^. All parents may benefit from heightened responsiveness to chick cues, but the neural strategies by which they achieve this may differ. Females, with complete certainty of maternity, might rely on more selective auditory tuning—enhancing responses to begging calls and, in some cases, specifically to their own offspring’s calls—to efficiently direct care toward their brood. Males, on the other hand, who have reduced paternity assurance, even in species with low extra-pair paternity rates^86^, may adopt a broader strategy by increasing auditory vigilance generally rather than forming narrowly tuned representations of particular chicks. Such a strategy could ensure that they respond robustly to any chick in the nest, regardless of relatedness, while maintaining the benefits of a stable monogamous partnership.

It is also possible that males rely more heavily on non-auditory components of the begging display, such as head-waggling movements, gape coloration patterns, or social cues from the female, which are known to influence parental provisioning decisions^87,88^. If males weigh these additional modalities more strongly, broad auditory enhancement may be sufficient without requiring fine auditory discrimination. These hypotheses remain speculative but provide testable predictions. Future work that compares auditory, visual, and multisensory cue use across sexes—and across different stages of chick development—could help clarify whether the observed sex differences in NCM reflect distinct parental strategies, differences in cue weighting, or other aspects of parental decision-making.

### Species differences in auditory processing of begging calls

Only one other study to date has looked at neural responses to chick stimuli in breeding and non-breeding birds. Specifically, Vidas-Guscic et al.^63^ used fMRI to measure brain activity in response to conspecific adult songs and nestling begging calls using wild-caught European starlings. Subjects were tested in a photostimulated state (mimicking breeding conditions) and again in a photorefractory state (mimicking on-breeding conditions). In agreement with our results, Vidas-Guscic et al. found that females and males in the breeding condition showed a heightened response to begging calls in NCM, relative to non-breeding birds, but not to other stimuli. In contrast to our study, however, they found birds in a non-breeding state *also* responded stronger to begging calls (compared to adult songs), whereas in our study we found that non-breeding zebra finches did not respond strongly to begging calls.

Although our two studies used different methods to measure neural activity, this likely cannot account for differences in results as both methods are able to detect increases and decreases in neural response strength to stimuli, albeit, at different time scales. In the Vidas-Guscic et al. study, birds were wild caught and brought into the lab for testing, and so their age and reproductive histories were not known. It is possible that these birds may have been reproductively experienced, and therefore, potentially exhibited increased baseline responses to begging calls due to prior reproductive experience. In addition, since all birds were tested in the same order (photostimulated/breeding state first, and photorefractory/non-breeding state second), all birds in the non-breeding state had already been exposed to chick begging stimuli in this context, and therefore, elevated baseline responses may reflect experience/memory of chick stimuli in this context, rather than overall higher baseline responsivity to begging calls.

Another possibility is that these results reflect species differences in baseline responses towards chick cues that are potentially driven by different ecological and/or evolutionary pressures. For example, while both species are socially monogamous, starlings have much higher rates of extra-pair fertilizations and mate-switching^89^, and therefore may favor broader sensitivity to chick cues, even in non-breeding adults. Running additional studies in multiple species comparing experiences and inexperienced breeders would help illuminate whether this is an effect of experience or some other species difference.

### Conclusions

In summary, our results demonstrate that auditory processing in the zebra finch NCM is dynamically shaped by parental state, revealing both state-dependent and sex-specific modulation of neural responses to offspring cues. Parenting birds exhibited enhanced responsiveness to chick begging calls, alongside elevated baseline firing in narrow-spiking (putative inhibitory) neurons, indicating a potentially broad reorganization of auditory circuit function during parental care. These effects may be supported by an interaction between neuroendocrine changes associated with parenting and experience-dependent plasticity arising from repeated exposure to chick vocalizations, but more work is needed in order to test these hypotheses. Although we did not observe population-level neural discrimination between own and novel chick calls at the age tested, emerging selectivity at the single-unit level—particularly in parenting females—suggests that individual recognition may develop later or be encoded in finer-scale neural dynamics. In addition, the distinct response profiles observed in males and females point to potentially different neural strategies for supporting convergent parental behaviors. Finally, comparison with European starlings’ neural responses to begging calls highlights that species-specific life histories, social systems, and ecological pressures may shape baseline sensitivity to chick cues in fundamentally different ways. Taken together, our findings establish NCM as a key site of state-dependent auditory plasticity during parenting and provide a neurophysiological framework for understanding how sensory systems adapt to meet the demands of parental care.

## Acknowledgements

We thank Yuni Gerzon, Pam Anderson, and Quincy Bartlett for assistance with brain sectioning, Matheus M. Macedo-Lima for advice and consulting on the electrophysiology methods, Michelle MacGray and other UMass animal care staff for their excellent animal husbandry, the members of the Remage-Healey lab for discussion and feedback on the data presented in this manuscript, and members of the Dantzer lab for feedback on earlier drafts of this paper.

## Data availability statement

The data that support the findings of this study are available from the corresponding author upon reasonable request.

## Supplemental figure

*Figure S1.* Each Y-axis row represents a single unit recording. The X-axis are time bins (50 ms) divided across the duration of the playback trial. Plotted colors are the average firing rates for each unit at a particular time bin. Warmer colors (reds) depict more selective responses, whereas cooler colors (blues) depict more unselective responses. White/neutral colors indicate no selectivity toward either stimulus.

## Supplemental Table

*Table S1.* Tukey post-hoc test for all combination of factors (sex, stimulus type, and parenting status) following the ANOVA used for the evoked NCM responses across treatment groups in Figure 2. Comparisons in italics is not shown on Figure 2 as they were not deemed to be biologically meaningful for the purposes of this study. P-values in red are significantly different (p<0.05). M=male, F=female, CI = confidence interval.

