## Supplementary figures and images for "Parental state dynamically reshapes auditory processing of offspring vocalizations in zebra finches"

### Supplemental Figure 1

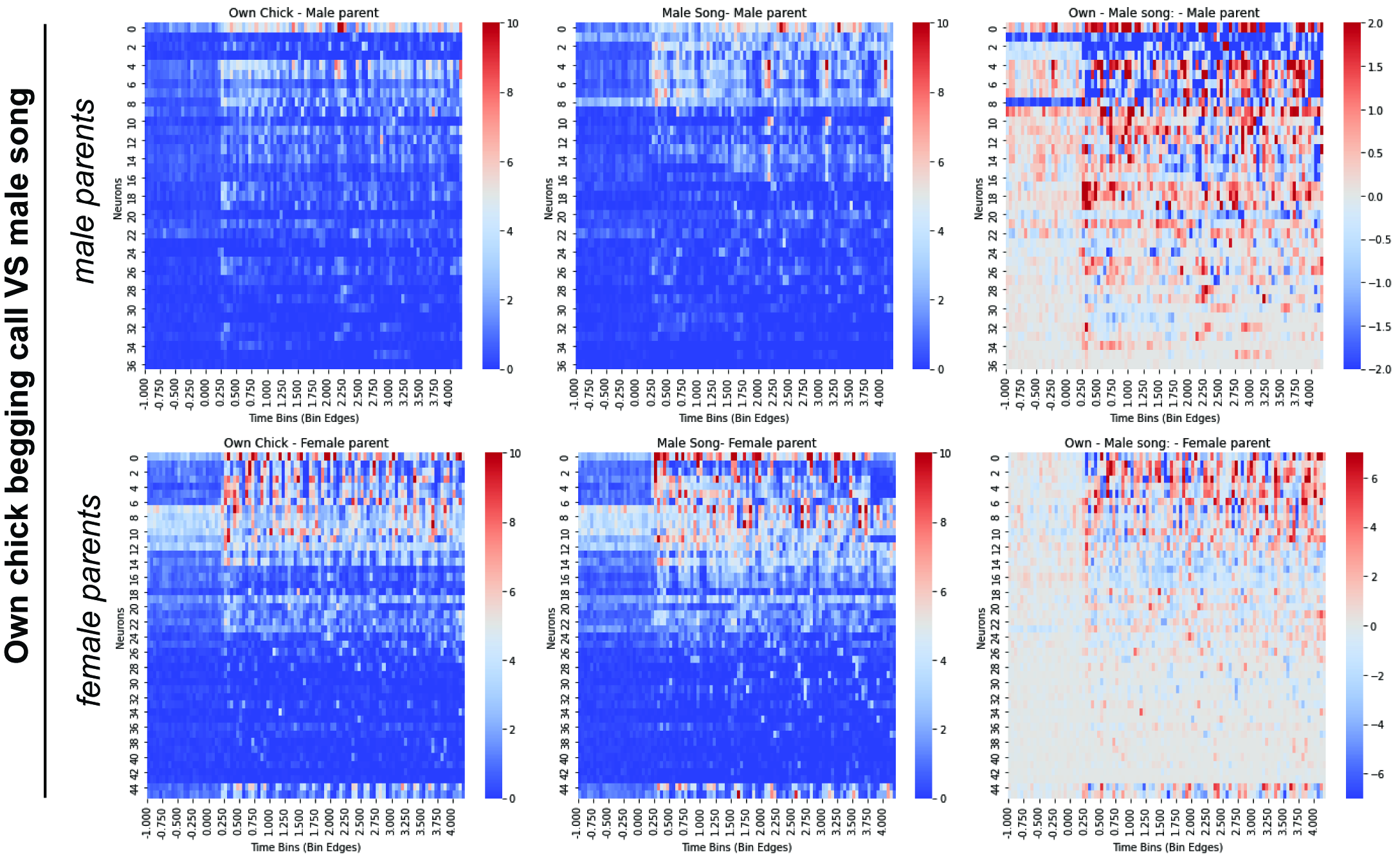
