## Supplemental Table 1 for "Parental state dynamically reshapes auditory processing of offspring vocalizations in zebra finches"

**Supplemental Table S1.**

| Group 1 | Group 2 | Mean difference | p-value | Lower CI | Upper CI |
|---:|---:|---:|---:|---:|---:|
| F & male_song & non-parenting | F & male_song & parent | 0.99 | 0.03 | 0.03 | 1.95 |
| F & male_song & non-parenting | F & novel_chick & non-parenting | -0.65 | 0.06 | -1.32 | 0.01 |
| F & male_song & non-parenting | F & novel_chick & parent | 0.38 | 0.99 | -0.58 | 1.34 |
| F & male_song & non-parenting | F & own_chick & parent | 1.01 | 0.09 | -0.07 | 2.09 |
| F & male_song & non-parenting | F & tone & non-parenting | -2.01 | 0.00 | -2.67 | -1.34 |
| *F & male_song & non-parenting* | *F & tone & parent* | *-1.56* | *0.00* | *-2.52* | *-0.60* |
| F & male_song & non-parenting | M & male_song & non-parenting | 1.01 | 0.83 | -0.77 | 2.79 |
| F & male_song & non-parenting | M & male_song & parent | 0.68 | 0.69 | -0.40 | 1.75 |
| F & male_song & non-parenting | M & novel_chick & non-parenting | -0.62 | 1.00 | -2.41 | 1.16 |
| F & male_song & non-parenting | M & novel_chick & parent | 0.77 | 0.49 | -0.31 | 1.84 |
| F & male_song & non-parenting | M & own_chick & parent | 0.77 | 0.48 | -0.31 | 1.85 |
| *F & male_song & non-parenting* | *M & tone & non-parenting* | *-2.24* | *0.00* | *-4.02* | *-0.46* |
| F & male_song & non-parenting | M & tone & parent | 0.03 | 1.00 | -1.05 | 1.10 |
| *F & male_song & parent* | *F & novel_chick & non-parenting* | *-1.64* | *0.00* | *-2.60* | *-0.69* |
| F & male_song & parent | F & novel_chick & parent | -0.61 | 0.90 | -1.79 | 0.57 |
| F & male_song & parent | F & own_chick & parent | 0.02 | 1.00 | -1.26 | 1.30 |
| *F & male_song & parent* | *F & tone & non-parenting* | *-3.00* | *0.00* | *-3.96* | *-2.04* |
| F & male_song & parent | F & tone & parent | -2.55 | 0.00 | -3.73 | -1.37 |
| F & male_song & parent | M & male_song & non-parenting | 0.02 | 1.00 | -1.89 | 1.93 |
| F & male_song & parent | M & male_song & parent | -0.31 | 1.00 | -1.59 | 0.97 |
| F & male_song & parent | M & novel_chick & non-parenting | -1.62 | 0.21 | -3.53 | 0.30 |
| F & male_song & parent | M & novel_chick & parent | -0.23 | 1.00 | -1.50 | 1.05 |
| F & male_song & parent | M & own_chick & parent | -0.22 | 1.00 | -1.50 | 1.06 |
| *F & male_song & parent* | *M & tone & non-parenting* | *-3.23* | *0.00* | *-5.14* | *-1.32* |

| | | | | | |
|---|---|---|---|---|---|
| F & male_song & parent | M & tone & parent | -0.97 | 0.38 | -2.24 | 0.31 |
| F & novel_chick & non-parenting | F & novel_chick & parent | 1.03 | 0.02 | 0.08 | 1.99 |
| F & novel_chick & non-parenting | F & own_chick & parent | 1.66 | 0.00 | 0.59 | 2.74 |
| F & novel_chick & non-parenting | F & tone & non-parenting | -1.35 | 0.00 | -2.02 | -0.69 |
| F & novel_chick & non-parenting | F & tone & parent | -0.91 | 0.09 | -1.86 | 0.05 |
| F & novel_chick & non-parenting | M & male_song & non-parenting | 1.66 | 0.10 | -0.12 | 3.44 |
| *F & novel_chick & non-parenting* | *M & male_song & parent* | *1.33* | *0.00* | *0.26* | *2.41* |
| F & novel_chick & non-parenting | M & novel_chick & non-parenting | 0.03 | 1.00 | -1.75 | 1.81 |
| *F & novel_chick & non-parenting* | *M & novel_chick & parent* | *1.42* | *0.00* | *0.34* | *2.49* |
| F & novel_chick & non-parenting | M & own_chick & parent | 1.42 | 0.00 | 0.35 | 2.50 |
| F & novel_chick & non-parenting | M & tone & non-parenting | -1.59 | 0.14 | -3.37 | 0.19 |
| F & novel_chick & non-parenting | M & tone & parent | 0.68 | 0.68 | -0.40 | 1.75 |
| F & novel_chick & parent | F & own_chick & parent | 0.63 | 0.93 | -0.65 | 1.91 |
| F & novel_chick & parent | F & tone & non-parenting | -2.39 | 0.00 | -3.35 | -1.43 |
| F & novel_chick & parent | F & tone & parent | -1.94 | 0.00 | -3.12 | -0.76 |
| F & novel_chick & parent | M & male_song & non-parenting | 0.63 | 1.00 | -1.28 | 2.54 |
| F & novel_chick & parent | M & male_song & parent | 0.30 | 1.00 | -0.98 | 1.58 |
| F & novel_chick & parent | M & novel_chick & non-parenting | -1.00 | 0.89 | -2.92 | 0.91 |
| F & novel_chick & parent | M & novel_chick & parent | 0.39 | 1.00 | -0.89 | 1.66 |
| F & novel_chick & parent | M & own_chick & parent | 0.39 | 1.00 | -0.89 | 1.67 |
| F & novel_chick & parent | M & tone & non-parenting | -2.62 | 0.00 | -4.53 | -0.71 |
| F & novel_chick & parent | M & tone & parent | -0.35 | 1.00 | -1.63 | 0.92 |
| F & own_chick & parent | F & tone & non-parenting | -3.02 | 0.00 | -4.09 | -1.94 |
| F & own_chick & parent | F & tone & parent | -2.57 | 0.00 | -3.85 | -1.29 |
| F & own_chick & parent | M & male_song & non-parenting | 0.00 | 1.00 | -1.97 | 1.97 |
| F & own_chick & parent | M & male_song & parent | -0.33 | 1.00 | -1.70 | 1.04 |
| F & own_chick & parent | M & novel_chick & non-parenting | -1.63 | 0.23 | -3.61 | 0.34 |
| F & own_chick & parent | M & novel_chick & parent | -0.24 | 1.00 | -1.61 | 1.12 |

| | | | | | |
|---|---|---|---|---|---|
| F & own_chick & parent | M & own_chick & parent | -0.24 | 1.00 | -1.61 | 1.13 |
| F & own_chick & parent | M & tone & non-parenting | -3.25 | 0.00 | -5.22 | -1.28 |
| F & own_chick & parent | M & tone & parent | -0.98 | 0.47 | -2.35 | 0.38 |
| F & tone & non-parenting | F & tone & parent | 0.45 | 0.95 | -0.51 | 1.41 |
| F & tone & non-parenting | M & male_song & non-parenting | 3.01 | 0.00 | 1.23 | 4.80 |
| F & tone & non-parenting | M & male_song & parent | 2.68 | 0.00 | 1.61 | 3.76 |
| F & tone & non-parenting | M & novel_chick & non-parenting | 1.38 | 0.34 | -0.40 | 3.16 |
| F & tone & non-parenting | M & novel_chick & parent | 2.77 | 0.00 | 1.70 | 3.85 |
| F & tone & non-parenting | M & own_chick & parent | 2.78 | 0.00 | 1.70 | 3.85 |
| F & tone & non-parenting | M & tone & non-parenting | -0.24 | 1.00 | -2.02 | 1.55 |
| F & tone & non-parenting | M & tone & parent | 2.03 | 0.00 | 0.96 | 3.11 |
| F & tone & parent | M & male_song & non-parenting | 2.57 | 0.00 | 0.66 | 4.48 |
| F & tone & parent | M & male_song & parent | 2.24 | 0.00 | 0.96 | 3.52 |
| F & tone & parent | M & novel_chick & non-parenting | 0.94 | 0.93 | -0.98 | 2.85 |
| F & tone & parent | M & novel_chick & parent | 2.33 | 0.00 | 1.05 | 3.60 |
| F & tone & parent | M & own_chick & parent | 2.33 | 0.00 | 1.05 | 3.61 |
| F & tone & parent | M & tone & non-parenting | -0.68 | 1.00 | -2.59 | 1.23 |
| F & tone & parent | M & tone & parent | 1.59 | 0.00 | 0.31 | 2.86 |
| M & male_song & non-parenting | M & male_song & parent | -0.33 | 1.00 | -2.30 | 1.64 |
| M & male_song & non-parenting | M & novel_chick & non-parenting | -1.63 | 0.59 | -4.06 | 0.80 |
| M & male_song & non-parenting | M & novel_chick & parent | -0.24 | 1.00 | -2.21 | 1.73 |
| M & male_song & non-parenting | M & own_chick & parent | -0.24 | 1.00 | -2.21 | 1.73 |
| M & male_song & non-parenting | M & tone & non-parenting | -3.25 | 0.00 | -5.68 | -0.82 |
| M & male_song & non-parenting | M & tone & parent | -0.98 | 0.92 | -2.95 | 0.99 |
| M & male_song & parent | M & novel_chick & non-parenting | -1.30 | 0.61 | -3.27 | 0.67 |
| M & male_song & parent | M & novel_chick & parent | 0.09 | 1.00 | -1.28 | 1.46 |
| M & male_song & parent | M & own_chick & parent | 0.09 | 1.00 | -1.28 | 1.46 |
| M & male_song & parent | M & tone & non-parenting | -2.92 | 0.00 | -4.89 | -0.95 |
| M & male_song & parent | M & tone & parent | -0.65 | 0.95 | -2.02 | 0.72 |
| M & novel_chick & non-parenting | M & novel_chick & parent | 1.39 | 0.50 | -0.58 | 3.36 |

| | | | | | |
|---|---|---|---|---|---|
| M & novel_chick & non-parenting | M & own_chick & parent | 1.39 | 0.50 | -0.58 | 3.37 |
| M & novel_chick & non-parenting | M & tone & non-parenting | -1.62 | 0.60 | -4.05 | 0.81 |
| M & novel_chick & non-parenting | M & tone & parent | 0.65 | 1.00 | -1.32 | 2.62 |
| M & novel_chick & parent | M & own_chick & parent | 0.00 | 1.00 | -1.36 | 1.37 |
| M & novel_chick & parent | M & tone & non-parenting | -3.01 | 0.00 | -4.98 | -1.04 |
| M & novel_chick & parent | M & tone & parent | -0.74 | 0.87 | -2.11 | 0.63 |
| M & own_chick & parent | M & tone & non-parenting | -3.01 | 0.00 | -4.98 | -1.04 |
| M & own_chick & parent | M & tone & parent | -0.74 | 0.86 | -2.11 | 0.62 |
| M & tone & non-parenting | M & tone & parent | 2.27 | 0.01 | 0.30 | 4.24 |
